# The Arabidopsis ARID-HMG protein AtHMGB15 modulates JA signalling by regulating MYC2 during pollen development

**DOI:** 10.1101/2022.11.10.515973

**Authors:** Sonal Sachdev, Ruby Biswas, Adrita Roy, Shubho Chaudhuri

## Abstract

In flowering plants, jasmonic acid (JA) signalling regulates the complex process of male gametophyte development. JA signalling initiates with the activation of MYC2 transcription factor, for the expression of several JA responsive genes throughout stamen development and pollen maturation. However, the regulation of JA signalling during different developmental stages of male gametophytes is still less understood. In this study we have characterized T-DNA insertion line of AtHMGB15. Phenotypic characterization of *athmgb15-4* mutant plants showed delayed bolting, shorter siliques and reduced seed set compared to wildtype. Moreover, deletion of AtHMGB15 resulted in defective pollen morphology, delayed pollen germination, abberant pollen tube growth and a higher percentage of non-viable pollen population in *athmgb15-4* compared to wildtype. Molecular analysis indicated down-regulation of JA-biosynthesis and JA-signalling genes *viz* MYC2, MYB21 and MYB24 in *athmgb15-4* mutant. Furthermore, jasmonic acid and its derivatives were found almost ten-fold lower in *athmgb15-4* flowers. However, exogenous application of jasmonate could restore pollen morphology and pollen germination, suggesting that impaired JA signalling is responsible for the pollen phenotype in *athmgb15* mutant. AtHMGB15 physically interacts with MYC2 protein to form the transcription activation complex for promoting transcription of genes responsible for JA signalling during stamen and pollen development. Collectively, our findings indicate that AtHMGB15, a plant specific DNA binding protein of the ARID-HMG group, acts as a positive regulator of JA signalling to control the spatiotemporal expression of key regulators responsible for stamen and pollen development.

## INTRODUCTION

The development of male gametophyte in angiosperms is a complex phenomenon that requires coordination of almost all major plant hormone signalling (Marciniak and Przedniczek, 2019, Mascarenhas, 1990, Wilson and Zhang, 2009). The spatiotemporal activity of key hormone signalling factors regulates the pollen maturation, anther dehiscence, release of pollen to the surface of stigma and pollen tube germination, for successful fertilization. Pollen development starts in anther with the differentiation of sporogenous cells (pollen mother cell) that undergo meiosis to form tetrads of haploid microspores. The development of free microspores starts with two rounds of mitotic divisions and the formation of pollen cell wall through programmed cell death (PCD) of the tapetum layer (McCormick, 2004, Zhang et al., 2007). Degeneration of tapetum layer is also important for anther dehiscence and release of the mature pollens. In self-pollinating plants, the release of mature pollen (anther dehiscence) on the surface of stigma depends upon the appropriate length of the stamen filament. The anthers in these self-pollinating plants are positioned at equivalent height or above the stigma papillae for efficient release of pollen and fertilization. Any defects during pollen maturation, stamen elongation or anther dehiscence can cause loss of fertility or complete male sterility.

Plant hormone jasmonic acid (JA) and its derivatives are indispensable for the development of stamen and male gametophyte maturation (Huang et al., 2017b). In Arabidopsis, JA-biosynthesis deficient mutants viz. *fad3fad7fad8, dad1, lox3-lox4, aos*, and *opr3* are male sterile due to arrested stamen development at anthesis (Caldelari et al., 2011, Ishiguro et al., 2001, McConn and Browse, 1996, Park et al., 2002, Stintzi and Browse, 2000). These mutants have indehiscent anthers or short filaments that fail to reach stigma surface. Although the pollens from these mutants develop normally to produce tricellular gametophyte but lost viability during later stages (Acosta and Przybyl, 2019). Exogenous application of jasmonic acid can restore the male sterile phenotype of JA-biosynthesis deficient mutants (Park et al., 2002). CORONATINE INSENSITIVE1 (COI1), a F-box protein is a part of SKP1-CULLIN1-F-box-type (SCF) E3 ubiquitin ligase complex SCF^COI1^, and an important component of JA signalling. COI1 form complex with transcriptional repressors JAZ in presence of JA-Ile derivative and ubiquitinate for 26S proteasome mediate degradation to release *MYC* transcription factor for JA responsive gene expression(Chini et al., 2007, Devoto et al., 2002, Thines et al., 2007, Xie et al., 1998, Zhai et al., 2015). Like JA-biosynthesis deficient mutants, *coi1* mutants are also impaired in stamen maturation and are male sterile however exogenous JA application cannot rescue *coi1* fertility (Feys et al., 1994, Xu et al., 2002).

The basic helix–loop–helix (bHLH) transcription factor *MYC2* is the key regulator of JA response. MYC2 activates the transcription by binding to G-box motif existing in the promoter regions of JA responsive genes (Dombrecht et al., 2007, Figueroa and Browse, 2012, Kazan and Manners, 2013, Pozo et al., 2008). In the absence or low concentration of JA-Ile, MYC2 activity was repressed by JAZ protein along with co-repressors TOPLESS (TPL), TPL-related (TPR) and adaptor protein NINJA (An et al., 2022, Huang et al., 2017a). An increase in JA-Ile concentration during development or environmental clues promotes the formation of COI-JA-JAZ co-receptor complex to promote COI-mediated degradation of JAZ through 26S proteosome to release MYC2 (Chini et al., 2009). Studies have shown that MYC2, MYC3, MYC4 and MYC5 function redundantly to regulate stamen development and seed production (Gao et al., 2016, Qi et al., 2015). While the single and double mutants showed no defect in stamen development; the triple mutants *myc2myc3myc4, myc2myc4myc5*, and *myc3myc4myc5* exhibited delayed stamen development (Dombrecht et al., 2007, Schweizer et al., 2013). The anthers of these triple mutants failed to dehisce at the floral stage13 and pollens were unable to germinate *in vitro*, however, anther dehiscence and pollen maturation occur at the later stage of flower development. The quadruple mutant in comparison to the triple mutant has more severe defects in stamen development with short stamen filament, indehiscent anther and nonviable pollens (Qi et al., 2015).

MYC coordinates JA signalling through R2R3 types of MYB transcription factors, MYB21 and MYB24 during stamen maturation (Song et al., 2011). MYB21 and MYB24 physically interact with MYC2 to form the MYC-MYB complex for transcription activation and interacts with JAZ to attenuate their activity (Yang et al., 2020, Zhang et al., 2021). The phytohormone gibberellin (GA) has been shown regulate the expression of MYB21/24 and promotes stamen growth (Cheng et al., 2009). Studies indicate that DELLA impedes JA biosynthesis by inhibiting the expression of *DAD1* and *LOX1*. DELLA also interacts with MYB21/24 in absence of GA and represses their transcriptional activity (Cheng et al., 2009). GA triggers the ubiquitination of DELLA, and upregulates the expression of JA biosynthesis gene *DAD1* and *LOX1* (Huang et al., 2020). The increased concentration of JA will induce the expression of *MYB21* and *MYB24* (Huang et al., 2020, Vera-Sirera et al., 2016). Thus, GA and JA signalling synergistically modulate stamen elongation by regulating MYC-MYB signalling (Chini et al., 2016, Song et al., 2014). *myb21* mutants have short filaments that unable the anthers to reach the pistil’s stigma resulting in complete male sterility (Mandaokar et al., 2006). However, *myb21* pollens are viable. *myb24* mutants are completely fertile, whereas *myb21myb24* double mutants are completely impaired stamen and are fully sterile suggesting that MYB21 alone is essential for filament elongation, while MYB24 promotes pollen viability and anther dehiscence(Huang et al., 2017a, Mandaokar and Browse, 2009, Mandaokar et al., 2006, Song et al., 2011).

AtHMGB15 belongs to a novel plant-specific HMG-box group of nuclear architectural proteins containing two DNA binding domain, ARID and HMG-box (Štros et al., 2007). Biochemical analysis shows that ARID-HMG proteins bind to different DNA topological structures preferably in the AT-rich region (Hansen et al., 2008, Roy et al., 2016). A previous study by Xia et.al demonstrated that AtHMGB15 plays an important role in pollen tube growth (Xia et al., 2014). Approximately 10% pollen grains of Ds insertion line of AtHMGB15 (*athmgb15-1*) have defective morphology. Comparative transcriptome between wildtype and *athmgb15-1* pollen showed alteration of genes specific for pollen. Further, it was shown that AtHMGB15 interact with two MIKC* transcription factors, AGL66 and AGL104. Although *athmgb15* mutant showed a defect in pollen development, it was not clear how AtHMGB15 contribute to this developmental process. With this background, we started characterising another mutant allele of *AtHMGB15 (athmgb15-4*) where the T-DNA was inserted at the first exon. Our study revealed that around 30% of pollens from *athmgb15-4* plants are defective in pollen morphology and most of the mutated pollens are round in shape with a defect in the reticulate pattern of ornamentation. Transcriptome analysis shows significant repression of JA biosynthesis and signalling in *athmgb15-4* flowers. Collectively our results indicated that AtHMGB15 regulates pollen development by regulating key master regulators of JA-signalling, MYC2, MYB21 and MYB24. This study is the first in-depth analysis to understand the mechanistic role of ARID-HMG protein in pollen development.

## RESULTS

### Isolation and Characterization of *athmgb15-4* mutant lines

The *athmgb15-4* mutant was screened from T-DNA insertion lines of *Arabidopsis* ecotype Col-0 from the GABI-Kat collection (GABI_351D08). GABI_351D08 from the GABI collection has the T-DNA insertion annotated at exon 1 of the gene At1g04880 (Fig 1A, i)). The T-DNA insertion contains sulfadiazine resistant marker. The homozygous *athmgb15-4* lines were obtained by self-crossing of heterozygous *athmgb15-4* plants followed by the selection of progeny showing sulfadiazine resistance. The homozygous lines were screened by PCR (Fig 1A,ii) and the T-DNA insertion was confirmed by Southern blot (Fig S1). q-RT-PCR analysis showed significant down-regulation of *AtHMGB15* expression in *athmgb15-4* mutant plants (Fig 1A,iii). We have shown previously the absence of AtHMGB15 protein in the same mutant (Mallik et al., 2020). The homozygous seeds were collected and used for subsequent studies.

**Figure 1:**
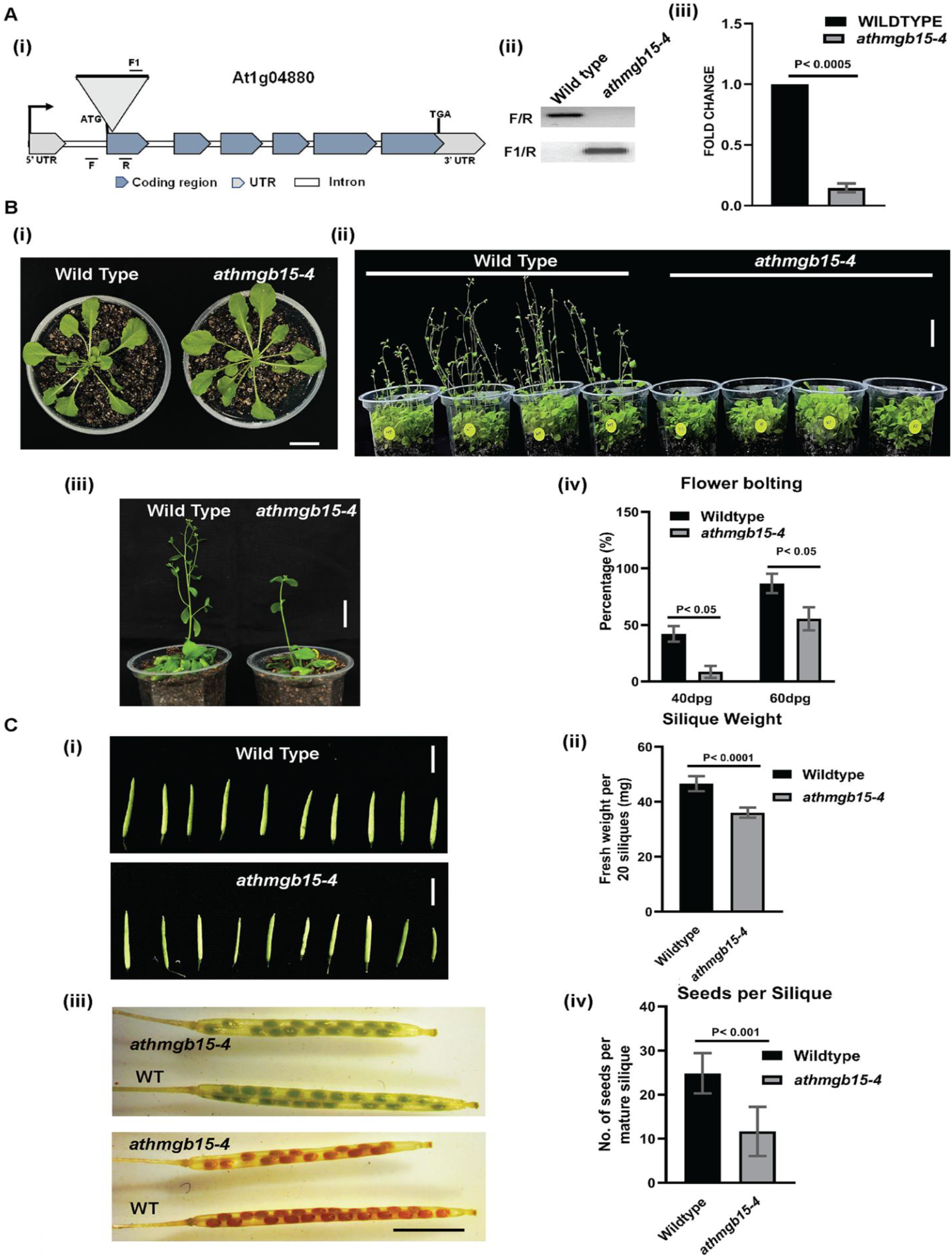
Phenotypic characterization of *athmgb15-4* mutant. **A (i)** Schematic showing the position of T-DNA insertion in the 1st exon of AtHMGB15 (At1g04880) and the position of PCR primers used for mutant screening. **(ii)** PCR confirmation of *athmgb15* homozygous line. **(iii)** q-RTPCR showing significant reduction of AtHMGB15 transcript in *athmgb15-4* lines. Error bars represent mean ± SD (n=3). **B (i)** Wild-type and *athmgb15-4* at the rosette stage. Scale bar=2cm. **(ii &iii)** Delayed flowering of *athmgb15-4* compared to wildtype. Scale bar=2cm. **(iv)** Quantitative analysis of flower bolting between *athmgb15-4* and wildtype. The experiments were done from seeds of 4-5 independent harvests. Data were collected from 100 plants of each batch and error bars represent mean ± SD (n=400) and significance was calculated by paired two-tailed student’s t-test (*denotes P ≤ 0.05). **C (i)** Comparative silique length of wildtype and *athmgb15-4*. Scale bar=5mm. **(ii)** quantitative silique fresh weight between athmgb15-4 and wildtype. Measurement was done using 20 siliques for each observation. Error bars represent mean ± SD (n=6). **(iii)** comparison of seed set between wildtype and *athmgb15-4*. Scale bar=2.5mm. **(iv)** seed numbers were counted from mature siliques of wildtype and *athmgb15-4*. Error bar represents mean ± SD (n=30). The significance of all these results was analysed by paired two-tailed student’s t-test.

The *athmgb15-4* mutant plants showed no phenotypic difference at the rosette stage (Fig 1B, i) except the primary root length *in athmgb15-4* appeared shorter compared to wild-type plants (Fig S2). Furthermore, in the flowering stage, *athmgb15-4* plants showed delayed bolting compared to wild type (Fig 1B, ii). Almost 45% (*p* ≤0.05) seedling of wildtype showed bolting after 40dpg compared to 8% (*p* ≤0.05) in *athmgb15-4* plants (Fig 1B, iii, iv). The seeds of *athmgb15-4* mutant plants showed no marked difference when compared with wild type. These mutant seeds are viable and germinate normally similar to wild type. However, mutant siliques were shorter in length compared to wild type (Fig 1C, i-ii) and had a lesser number of fertilized ovules resulting in less seed yield compared to wildtype plants (Fig 1C, iii-iv). Some of these observed phenotypes of *athmgb15-4* agree with previous observations reported by Xia *et al* using another mutant allele of *AtHMGB15* (Xia et al., 2014).

### AtHMGB15 mutation causes a defect in pollen morphology and delayed pollen germination rate

An earlier report has shown that *athmgb1*5 mutant (*athmgb15-1*) plants have defective pollen morphology (~10%) (Xia et al., 2014). Scanning electron microscopy (SEM) analysis revealed that the wild-type pollens are ellipsoidal (Fig 2A) whereas pollens of *athmgb15-4* plants have mixed shaped (Fig S3). While ellipsoid-shaped pollens were observed in the mutant pollen population, we have also observed around 25-30% of pollens having a circular shape and sometimes completely irregular shape (Fig 2B). Additionally, the outermost exine wall of wildtype pollens has a typical reticulate pattern of ornamentation, which is completely absent in the defective pollens of mutant pollen plants. To further understand the molecular changes in *athmgb15-4* plants that regulates pollen development, a comparative transcriptome approach was taken. Arabidopsis flowers (stage 13) were collected from wild-type and *athmgb15-4* plants and total RNA isolated from three independent sets was pooled and subjected to RNA-sequencing using the Illumina platform. Analysis of RNA-seq data showed significant down-regulation of genes involved in cell wall biosynthesis in *athmgb15-4* flowers (Fig 2C and Fig S4). Some of these cell wall genes includes pectin lyase, cellulose synthase, pectin methylesterase, extensin which are known to be involved in pollen development.

**Figure 2:**
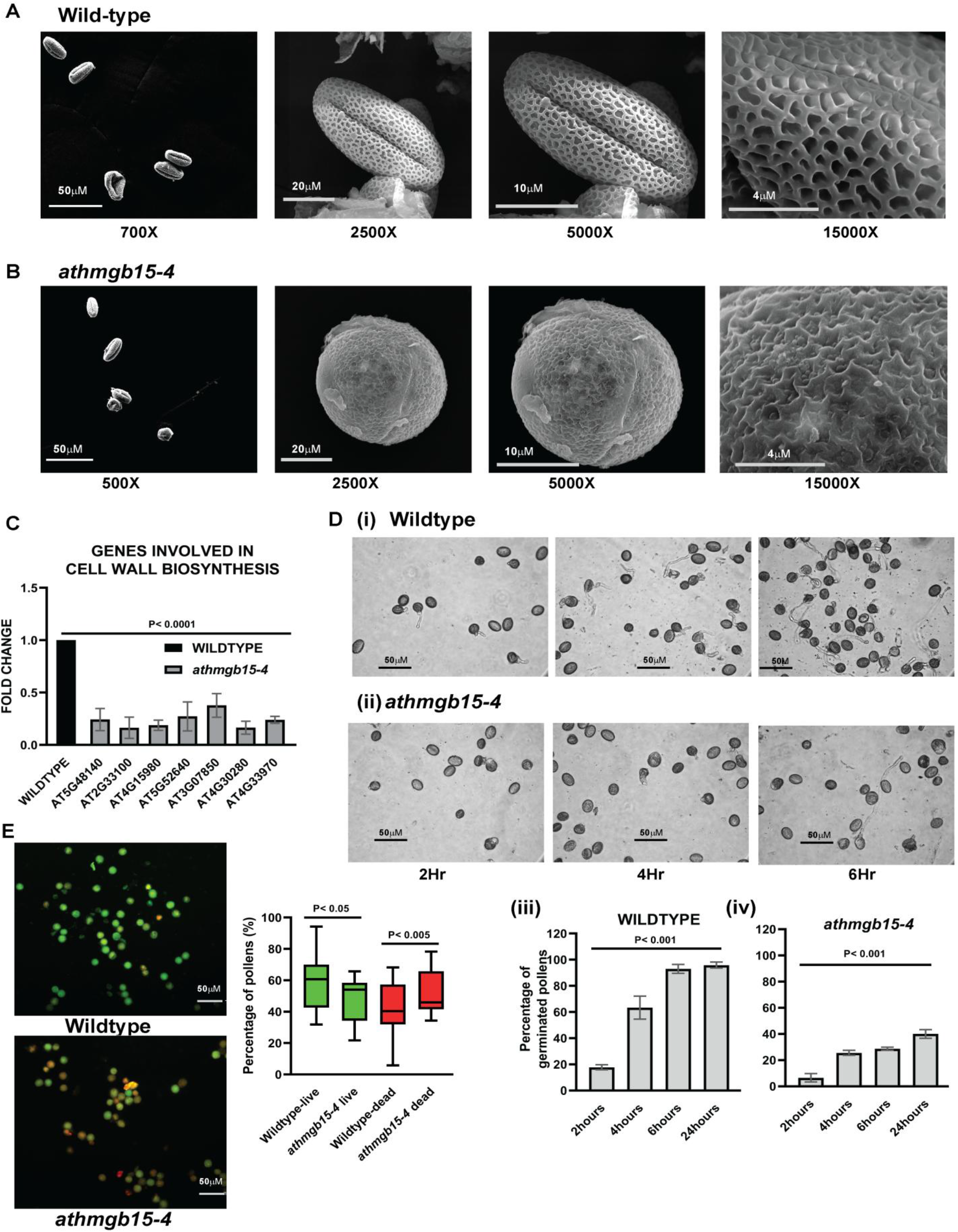
*athmgb15-4* have impaired pollen morphology and pollen germination compared to wildtype. **A.** Scanning electron microscopy (SEM) of pollens isolate from wildtype showing ellipsoidal shape with reticulate ornamentations. **B**. Representation of defective pollen morphology of *athmgb15-4* mutant having a circular shape with irregular ornamentation. The experiment was repeated at least 10 times with pollens isolated from different batches of wildytype and *athmgb15-4*. **C**. Expression of cell wall biosynthesis genes between wildtype and *athmgb15-4* using q-RTPCR. The fold change was represented with respect to wildtype. Error bars represent mean ± SD (n=3) and significance (*p*≤0.0005) was analysed by paired two-tailed student’s t-test. **D.** Freshly isolated pollens from **(i)** wildtype and **(ii)** *athmgb15-4* were subjected to *in vitro* germination for different time periods. **(iii) & (iv)** Graphical representation of rate of pollen germination of wildtype and *athmgb15-4* respectively. Error bars represent mean ± SD (n=5) and the significance of the result was analysed by one-way ANOVA (*p*≤0.005). **E**. Pollen viability was measured using fluorescein diacetate and propidium iodide. Box plot representation of pollen viability between wildtype and *athmgb15-4*. Error bars represent mean ± SD (n=12). The significance of all these results was analysed by paired two-tailed student’s t-test.

We subsequently examined the pollen germination rate between wildtype and *athmgb15-4* pollens. The time kinetics of *in vitro* pollen tube germination shows that within 4hrs, more than 50% of pollens (*p* ≤0.005) were germinated and by 6hrs almost 80% (*p* ≤0.005) germination was achieved for wildtype pollens (Fig 2D, i, iii). Interestingly, 40% (*p* ≤0.005) germination of *athmgb15-4* pollens was observed after 24hrs in pollen germination media (Fig 2D, ii, iv). These results indicate that mutation of *AtHMGB15* gene causes a severe defect in pollen morphology and significant delay in pollen tube germination rate. Note that some of the healthy pollens of *athmgb15-4* that start germinating from the beginning showed similar pollen tube length as of wildtype, although the percentage of fully germinating pollens was very low in mutants. The next obvious question that arises from these observations is whether the *athmgb15-4* pollens are viable. To answer this question, we isolated the wildtype and *athmgb15-4* pollens and stained them FDA (fluorescein diacetate) and PI (Propidium Iodide). While FDA is permeable to the cell membrane and can stain live cells, PI is impermeable and can stain DNA only when the cell integrity is compromised. Thus, PI-stained cells are considered dead cells. Comparison of differential staining of pollens with FDA and PI showed that a higher percentage of non-viable pollens (55%, *p*≤0.005) in *athmgb15-4* plants compared to wild-type (30%,*p* ≤0.05); thereby, justifying a lower number of germinating pollen population in mutant plants (Fig 2E).

### Deletion of *AtHMGB15* causes down-regulation of the jasmonic acid pathway during flower development

KEGG analysis of RNA-seq data shows enrichment of α-linolenic acid metabolism pathway, associated with differential gene expression (Fig. 3A, i). α-linolenic acid is the precursor of the plant phytohormone, jasmonic acid. Jasmonic acid and its derivates have been shown to regulate many developmental processes including stamen development and flowering (Jang et al., 2020, Wasternack and Hause, 2013). Interestingly, function annotation clustering using DAVID software (v 6.8) shows enrichment of Jasmonic acid (JA) biosynthesis and signalling with the data set (Fig 3A, ii). Further, the heatmap analysis constructed using genes involved in JA biosynthesis and signalling pathway displayed down-regulation of gene expression, in *athmgb15-4* flowers compared to wild-type (Fig. 3B). The expression of many JA biosynthesis and signalling genes were further validated using q-RTPCR. As shown in figure 3C, the relative fold change for genes from JA biosynthesis and JA signalling were significantly down-regulated in *athmgb15-4* flowers compared to wild-type; thus, validating our RNA-seq data.

**Figure 3:**
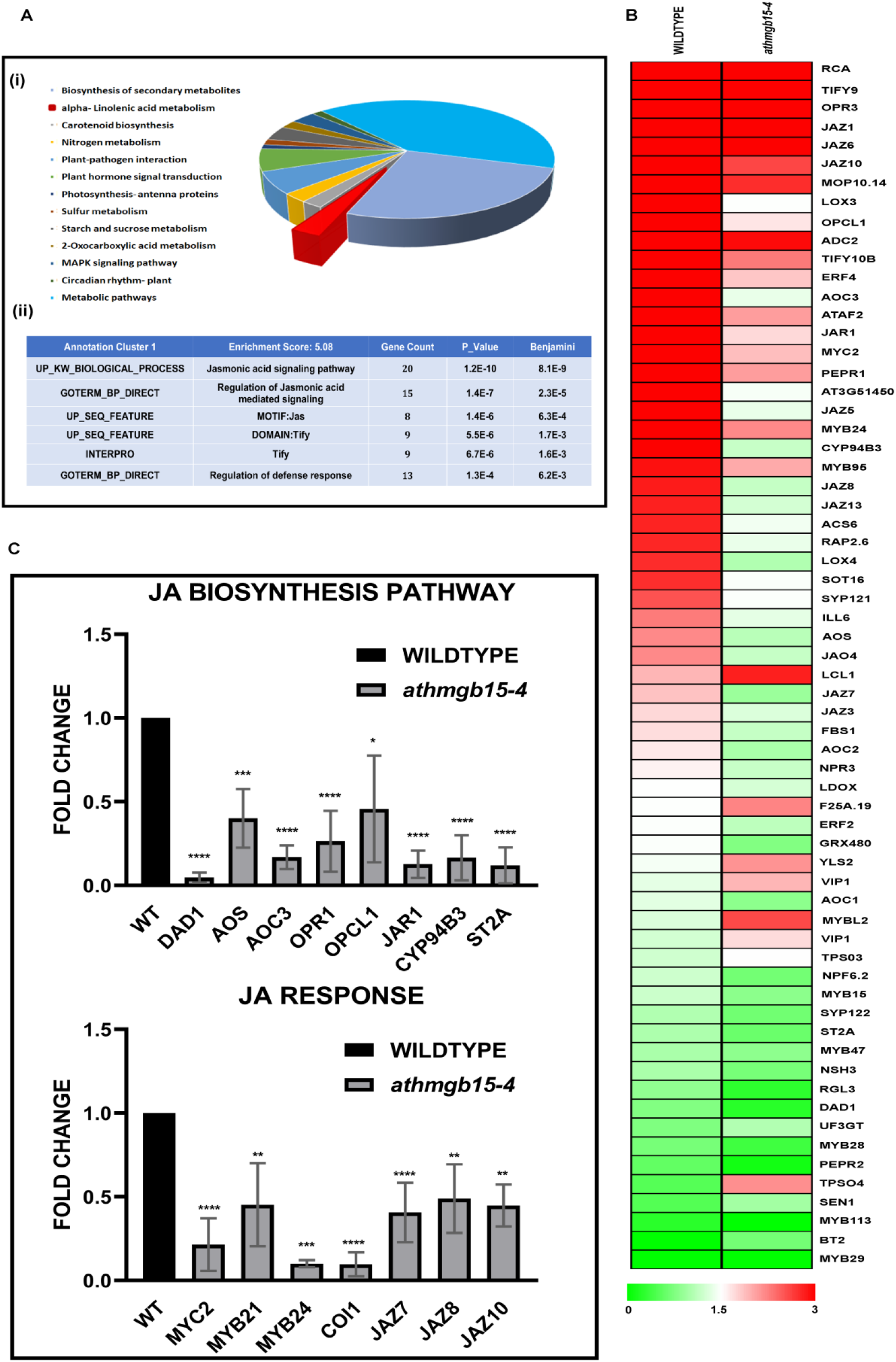
AtHMGB15 deletion affects jasmonic acid pathway. Comparative transcriptome between wildtype and *athmgb15* flowers were performed to identify the candidate genes involved in pollen development. **A (i)** KEGG analysis for significant DEGs using DAVID v 6.8 **(ii)** Functional annotation clustering showing enrichment of jasmonic acid pathway. **B**. Heatmap generated with log10(FPKM) of genes involved in JA biosynthesis and signalling. **C**. Expression of differentially regulated JA biosynthesis and signalling genes was analysed between wildtype and *athmgb15-4* using q-RTPCR. The fold change was represented with respect to wildtype. Error bars represent mean ± SD (n=3). The significance of all these results was analysed by paired two-tailed Student’s t-test. * Denotes p≤0.05, **p≤0.005, ***p≤0.0005 and **** p≤0.00005.

To further establish the role of *AtHMGB15* in JA signalling during pollen development, we then raised complementation lines in *athmgb15-4* background using full-length *AtHMGB15* gene under 35S constitutive promoter. Stable homozygous lines were selected (*athmgb15-4-OEA4*) and the expression of *AtHMGB15* was analysed using qRT-PCR (Fig 4A,iii). These complementation lines were found to be stable and recovered delayed bolting and small silique size phenotype of *athmgb15-4* mutant (Fig 4A, i, ii). A comparison of pollen tube germination rate between the complementation line and *athmgb15-4* showed a significantly higher population of germinated pollens in the complementation line compared to *athmgb15-4* (Fig 4B). The complementation line has more than 95% (*p* ≤0.05) of pollen in ellipsoidal shape indicating that the pollen morphology is completely recovered in these lines (Fig 4C). qRT-PCR results revealed higher expression of JA biosynthesis and signalling genes in the complementation line compared to the mutant (Fig 4D). The expression of JA biosynthesis genes in the complementation line was comparable to wild type however the expression of JA signalling genes was higher than wild type. Molecular and phenotypic analysis of complementation lines strongly indicates the role of *AtHMGB15* in JA mediated signalling events during pollen development.

**Figure 4:**
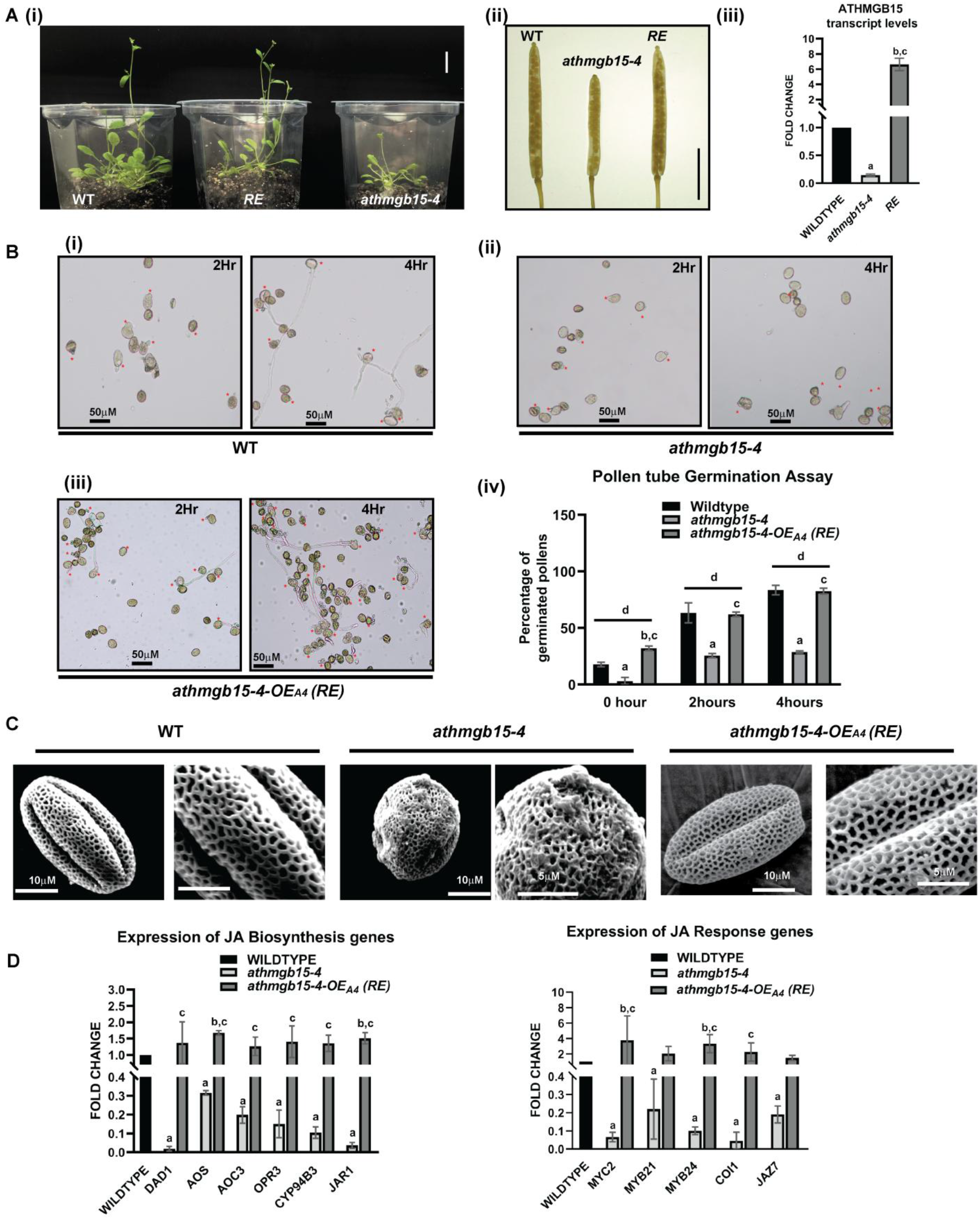
Complementation of *athmgb15-4* mutant line with AtHMGB15 restores pollen morphology and pollen tube germination. **A (i)** comparative flower bolting between wildtype, *athmgb15-4* and *athmgb15-4-OEA4* (RE). Scale bar=2cm. **(ii)** silique length of wildtype, *athmgb15-4* and RE. Scale bar=4mm. **(iii)** qRT-PCR to check AtHMGB15 transcript level in wildtype, *athmgb15-4* and RE. The fold change was represented with respect to wildtype. Error bars represent mean ± SD (n=3) and statistical significance (*p*≤0.05) was analysed by paired two-tailed student’s t-test. **a** denotes a significant difference between wildtype vs *athmgb15-4*, **b** denotes between wildtype vs RE, **c** denotes between *athmgb15-4* vs RE. **B. (i-iii)** comparative *in vitro* pollen germination between wildtype, *athmgb15-4* and RE **(iv)** quantification of the rate of pollen germination. Error bars represent mean ± SD (n=3). Statistical significance (p≤0.05) was analysed by two-way ANOVA with Fisher’s LSD test, **a** denotes a significant difference between wildtype vs *athmgb15-4*, **b** denotes between wildtype vs RE, **c** denotes between *athmgb15-4* vs RE and **d** denotes the significance of the three samples within the time point. **C.** SEM analysis of pollen morphology. **D** expression of JA biosynthesis and signalling genes in wildtype, *athmgb15-4* and RE flowers. The fold change was represented with respect to wildtype. Error bars represent mean ± SD (n=3) and significance (*p*≤0.05) was analysed by paired two-tailed student’s t-test. **a** denotes a significant difference between wildtype vs *athmgb15-4*, **b** denotes between wildtype vs RE, **c** denotes between *athmgb15-4* vs RE.

### *athmgb15* flowers show low levels of jasmonic acid and its derivatives

The down-regulation of JA biosynthesis genes in *athmgb15-4* mutants suggests a low intrinsic level of jasmonic acid and its derivatives. To check the hormone level, we next estimated the *in vivo* level of jasmonate in the flowers of wild-type, *athmgb15-4* and the complementation line *athmgb15-4*-OEA4. As shown in figure 5A, the levels of JA along with two of its derivatives methyl-jasmonate (MeJA) and JA-isoleucine (JA-Ile) are almost ten-fold (*p* ≤0.05) lower in *athmgb15* flowers compared to wild type. Furthermore, in the complementation line, the level of JA and its derivatives increased compared to *athmgb15-4*.

**Figure 5:**
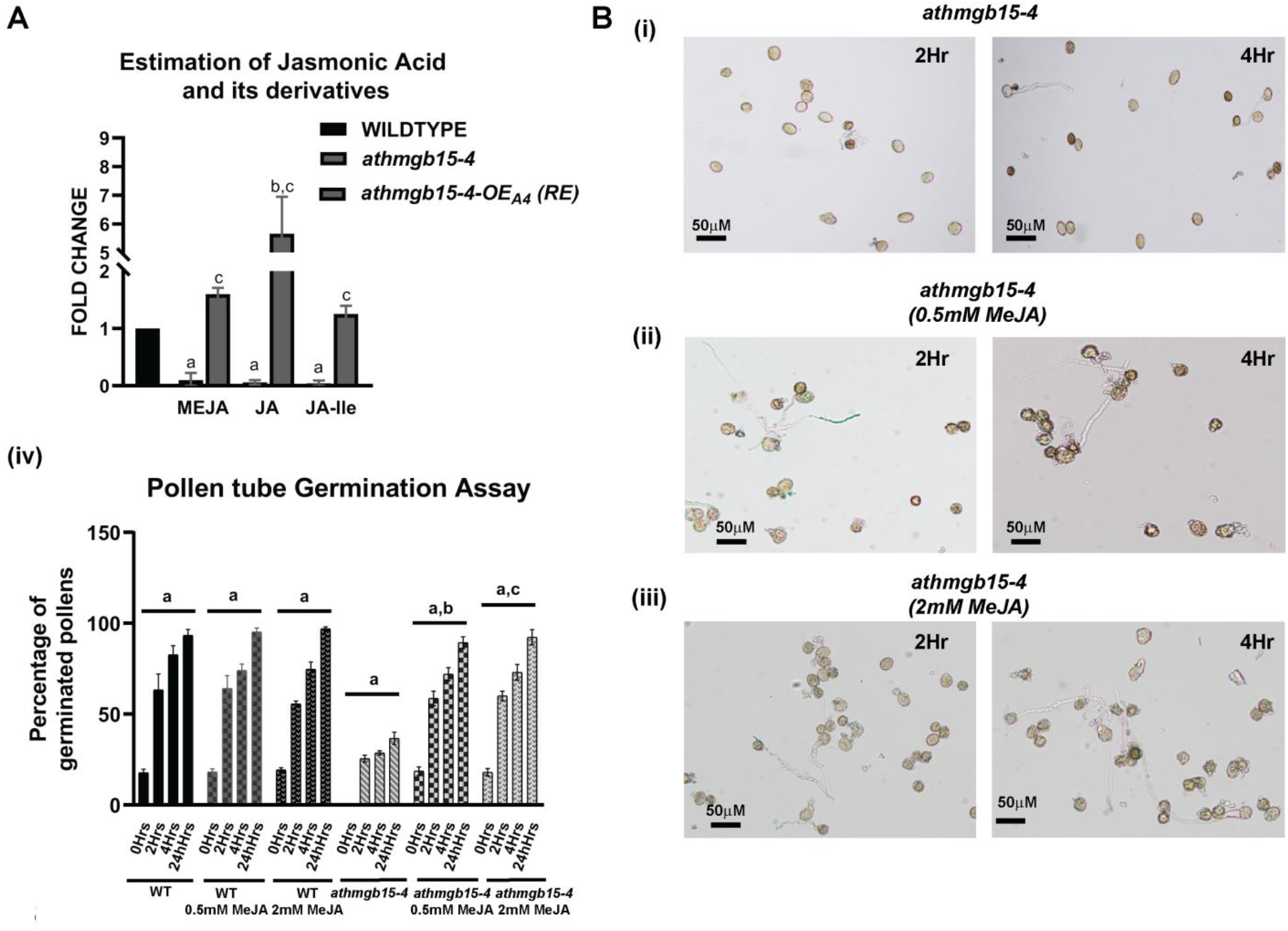
*athmgb15-4* mutants have reduced levels of JA and its derivatives. **A**. JA and its derivatives were measured from the flowers of wildtype, *athmgb15-4* and RE and represented as fold change with respect to wildtype. Error bars represent mean ± SD (n=3) with significance (p≤0.05) was analysed by paired two-tailed student’s t-test. **a** denotes a significant difference between wildtype vs *athmgb15-4*, **b** denotes between wildtype vs RE and c between *athmgb15-4* vs RE. **B. (i-iii)** Restoration of in vitro pollen germination of *athmgb15-4* on treatment with exogenous methyl jasmonate (0.5mM and 2mM). **(iv)** quantification of the rate of pollen tube germination in presence of different concentrations of methyl jasmonate. Error bars represent mean ± SD (n=3). Statistical significance was analysed by two-way ANOVA with Fisher’s LSD (p≤0.05). **a** denotes significance within the samples, **b** denotes significance between *athmgb15-4* vs *athmgb15-4* with 0.5mM MeJA and **c** denotes significance between *athmgb15-4* vs *athmgb15-4* with 2mM MeJA.

Since *athmgb15-4* flowers have low jasmonic acid, we studied the effect of exogenous JA application on *athmgb15-4* flowers by examining the pollen tube germination post 48hrs treatment. The result shows that exogenous treatment of methyl-jasmonate restores the pollen tube germination of *athmgb15-4* pollens, and the rate is equivalent to that of wild-type pollens (Fig 5B).

### AtHMGB15 acts as a transcription activator for the expression of *MYC2*

The transcriptome data and q-RTPCR results indicate down-regulation of the key transcription factors of JA-signalling *viz MYC2*, *MYB21* and *MYB24*, in *athmgb15-4* mutant flowers. This observation prompted us to check whether AtHMGB15 acts as a transcription activator for the expression of these genes.

#### AtHMGB15 occupancy at the upstream region of MYC2, MYB21 and MYB24

We performed ChIP assay using AtHMGB15 antibody and the immunoprecipitated DNA was subjected to q-PCR. The data were normalized with two loci, At1g01840 and At1g01310, showing no AtHMGB15 occupancy from our previous study (Mallik et al., 2020). The primers were designed from *in silico* analysis of ~2Kb promoter/upstream fragments that contain previously identified AtHMGB15 binding site A(A/C)--ATA---(A/T)(A/T) (Mallik et al., 2020). The q-PCR analysis showed AtHMGB15 occupancy at the promoter/upstream region of *MYC2, MYB21* as well as *MYB24* (Fig 6A). To test whether AtHMGB15 directly binds to the promoter/upstream region of *MYC2, MYB21* and *MYB24*, we performed *in vitro* DNA binding using purified recombinant AtHMGB15 protein. The EMSA results confirmed the direct binding of AtHMGB15 protein at the promoter regions of *MYC2, MYB21* and *MYB24* (Fig 6B).

**Figure 6:**
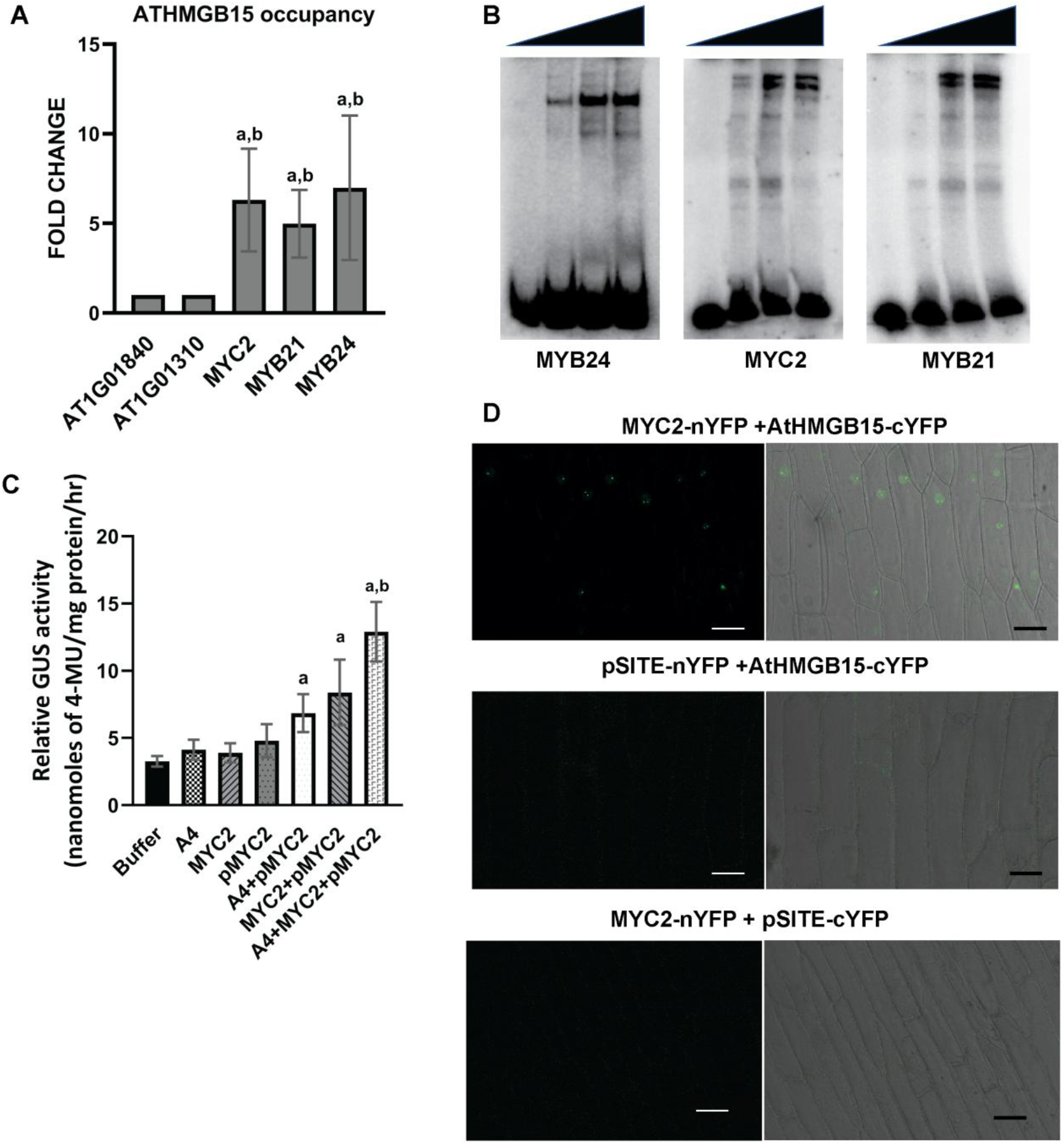
AtHMGB15 acts as a transcriptional activator for the expression of *MYC2*. **A**. ChIP analysis shows AtHMGB15 occupancy at the promoter/ upstream of *MYC2, MYB21* and *MYB24*. The data was normalised with no binding regions corresponding to At1g01840 and At1g01310. Error bars represent mean ± SD (n=3). The significance of the result was analysed by paired two-tailed student’s t-test (p≤0.05). **a** denotes significance when normalized with At1g01840 and **b** denotes normalized with At1g01310. **B**. EMSA showing binding of recombinant AtHMGB15 to 32P labelled DNA fragments correspond to the upstream region of *MYC2, MYB21* and *MYB24*. **C**. 2kb promoter region of MYC2 (pMYC2) was cloned with GUS reporter and Agrobacterium mediated infiltration was done with 35S::AtHMGB15 and 35S::MYC2 in *Nicotiana tabacum*. GUS reporter gene assay was done after 48hrs using MUG. Error bars represent mean ± SD (n=15). Statistical significance was analysed by paired two-tailed Student’s t-test (p≤0.05). **a** denotes significance between pMYC2 and pMYC2 with different combination of proteins used in the experiment and **b** denotes significance between pMYC2+AtHMGB15(A4) and pMYC2+AtHMGB15 (A4) + MYC2 proteins. **D**. BiFC confirming the interaction between AtHMGB15 and MYC2 in onion epidermal cells using split YPF. AtHMGB15-cYFP +pSITE-nYFP-C1 and MYC2-nYFP +pSITE-cYFP-N1 was used as control. Scale bar=50μm.

#### AtHMGB15 activates the transcription of MYC2

The binding of AtHMGB15 at the promoter/upstream region of *MYC2* prompts us to investigate whether AtHMGB15 regulates the transcription of *MYC2*. For this, ~2Kb promoter/upstream region of *MYC2* was cloned against GUS reporter gene in pCambia1304 replacing 35S promoter.

These constructs were infiltered into tobacco plants to examine the promoter activity in the absence and presence of AtHMGB15. AtHMGB15 is not a transcription factor but it can modulate transcription when associated with a transcription factor. Previous studies have identified the MYC2 binding site at the promoter of *MYC2* (Zander et al., 2020). Thus, we presumed that probably, AtHMGB15 can act as the co-activator of the transcription factor MYC2. To prove this hypothesis, we measured the promoter activity of *MYC2* in presence of both the proteins, MYC2 and AtHMGB15. As shown in figure 6C, the promoter activity of *MYC2* increases in presence of AtHMGB15 (pMYC2 +A4) as compared to only promoter (pMYC2). The increase was more with MYC2 (pMYC2+MYC2); supporting the earlier finding that MYC2 regulates its own transcription. Furthermore, in presence of both MYC2 and AtHMGB15, the promoter activity of pMYC2 was significantly higher to pMYC2. The result suggests that AtHMGB15 along with MYC2 TF positively activates the transcription of *MYC2*.

#### AtHMGB15 interacts with MYC2 protein to form the activator complex

Since AtHMGB15 along with MYC2 activates the transcription of pMYC2, we were interested to see whether they physically interact *in vivo* to form the activator complex. *AtHMGB15* and *MYC2* coding sequences were cloned in pSITE-cYFP-N1 and pSITE-nYFP-C1 respectively and used for BiFC using Agrobacterium mediated co-infiltration in onion epidermis. As shown in Fig 6D, AtHMGB15 interacts with MYC2 protein in the nucleus, particularly in the nucleolus. There was no YFP fluorescence observed in control combinations.

### AtHMGB15 promotes the transcription of *MYBs*

MYC2 has been shown to interact with R2R3-MYB transcription factors, MYB21 and MYB24 to regulate anther and pollen development in a JA dependent manner (Goossens et al., 2017). Our results confirm that the expression of *MYB21* and *MYB24* is down-regulated in *athmgb15* mutants. Also, the *in-silico* analysis shows the presence of MYC2 binding sites at the promoter region of *MYB24*.

Since AtHMGB15 and MYC2 proteins act as transcription activator complex, it was interesting to study whether this complex regulates the transcription of R2R3-MYBs transcription factors. To test this possibility, we first checked the expression of MYB21, MYB24 in the flowers of two previously characterised *myc2* knockout lines *myc2-2* and *jin1-2* (Boter et al., 2004, Lorenzo et al., 2004). The results show that expression of *MYB21* and *MYB24* were significantly downregulated in *myc2* mutant lines (Fig7A, i). The expression of JA biosynthesis gene *OPR3*, which was previously shown to be MYC2 dependent (Mandaokar et al., 2006), was found downregulated in these mutants. Subsequently, we have analysed JA content of myc2 mutant and found significant down-regulation of jasmonate contents in these two mutants (Fig 7A, ii). Interestingly, previous study has shown down-regulation of *MYB21* and *MYB24* expression in opr3 mutant and it can be restored by the application of exogenous JA (Mandaokar et al., 2006).

**Figure 7:**
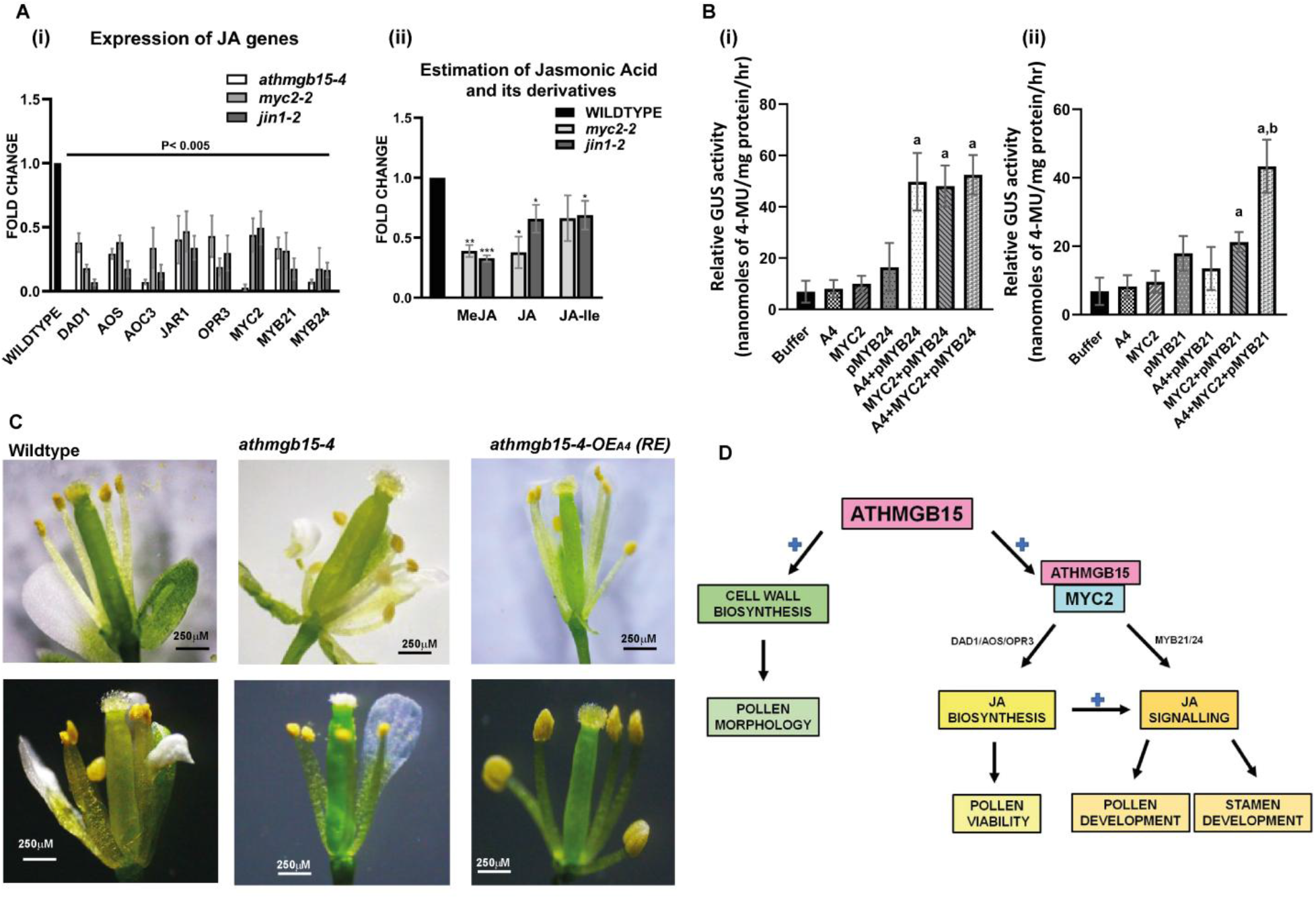
AtHMGB15 promotes transcription of *MYBs*. **A. (i)** Expression of JA genes in *MYC2* knockout mutants *myc2-2* and *jin1-2* and compared with the expression in *athmgb15-4* using q-RTPCR. The fold change was represented with respect to wildtype. Error bars represent mean ± SD (n=4). and the significance of the result was analysed by one-way ANOVA with Fisher’s LSD (p≤0.005). **(ii)** comparative JA and its derivatives content in flowers of wildtype and *MYC2* knockout mutants *myc2-2* and *jin1-2*. Error bars represent mean ± SD (n=3). The significance of all these results was analysed by paired two-tailed Student’s t-test. * Denotes p≤0.05, **p≤0.005 and ***p≤0.0005. **B**. **(i & ii)** 2kb promoter regions of MYB21 (pMYB21) and MYB24 (pMYB24) were cloned with GUS reporter and Agrobacterium mediated infiltration was done with 35S::AtHMGB15 and 35S::MYC2 in *Nicotiana tabacum*. GUS reporter gene assay was done after 48hrs using MUG. Error bars represent mean ± SD (n=15) and significance was analysed by paired two-tailed Student’s t-test (p≤0.05). **a** denotes significance between pMYB24/pMYB21 with different combination of proteins used in the experiment and **b** denotes significance between pMYB21+AtHMGB15(A4)/pMYB21+MYC2 and pMYB21+AtHMGB15 (A4) +MYC2 proteins **C**. Comparison of stamen phenotype between wildtype, *athmgb15* and *athmgb15-4-OEA4* (RE). **D.** Proposed model elucidating the role of AtHMGB15 in activating the JA pathways by forming an activation complex with MYC2 to regulate stamen and pollen development.

We next analysed the promoter activity of *MYB21* and *MYB24* in presence of AtHMGB15 and MYC2 transcription factor. The promoter activity of pMYB24 was significantly upregulated in presence of AtHMGB15 and MYC2 independently, however, in presence of both proteins, there was no additional increase in promoter activity (Fig 7B, i). For pMYB21, an increase in promoter activity was observed only in the presence of MYC2 protein (Fig 7B, ii,). There was no increase in the promoter activity in presence of AtHMGB15, although strong DNA binding activity of AtHMGB15 was observed in the promoter region. Interestingly, the activity of pMYB21 increase significantly in presence of AtHMGB15 and MYC2, suggesting that these two proteins form the activation complex for activating pMYB21.

Since R2R3 transcription factors were needed for the elongation of stamen during flower development, we further analysed the flower morphology of *athmgb15-4* and compared it to wildtype Arabidopsis flower. Our observation suggests that around 30% of *athmgb15-4* flowers have shorter stamen filaments compared to wild-type (Fig 7C, S5). This may be one of the reasons for poor fertilization and low seed yield in *athmgb15* mutants. The complementation lines on the other hand showed a similar stamen phenotype as compared to wildtype. Collectively, these results indicate that AtHMGB15 regulates the transcription of R2R3-MYB transcription factors during flower development to regulate the growth and development of stamen and pollens.

## DISCUSSION

### Deletion of AtHMGB15 impairs pollen morphology in *Arabidopsis*

The development of functional gametes and their respective floral organs is necessary for maximum pollination and genetic diversity. These complex processes are precisely regulated by endogenous cues. In this study, we have investigated the role of Arabidopsis ARID/HMG group of transcriptional regulators, AtHMGB15, in pollen development. Previous study by Xia *et.al*. using a Ds insertion line (Xia et al., 2014) and our study using another T-DNA mutant allele of AtHMGB15 showed defective pollen morphology and retarded pollen growth in mutant plants. These allelic mutants of *athmgb15* have a significant reduction in seed set. Since deletion of functional AtHMGB15 causes defective pollen morphology and pollen viability, we thought of investigating its role in pollen formation and maturation stages of floral development in *Arabidopsis*. Pollen development starts post meiosis of sporogenous cells that corresponds to stage 10 of floral development and continues through stage 12-13 till the completion of the pollen cell wall (Sanders et al., 1999). This is followed by elongation of filament and anther dehiscence to release viable pollens for germination (Goldberg et al., 1993, Scott et al., 2004). With this idea, we first investigated differential gene expression between wildtype and *athmgb15* mutant during the early stages of flower development, to identify AtHMGB15 regulated targets involved in the pollen development process. Our results indicated that some of genes responsible for pollen cell wall development were down-regulated in *athmgb15* mutant. Down-regulation of cell wall genes may explain the defective morphology and deformed cell wall architecture of *athmgb15* pollens.

Transcriptome analysis of wildtype and *athmgb15* revealed that jasmonic acid biosynthesis and response pathway are significantly downregulated in the mutant flowers. Downregulation of JA biosynthesis genes causes a significant decrease in jasmonate level in *athmgb15* mutant. Furthermore, the complementation of *athmgb15-4* with full-length AtHMGB15 or exogenous application of jasmonate completely restores impaired JA signalling and pollen morphology of *athmgb15-4*. AtHMGB15 mediated regulation of JA signalling explains the defective pollen development in *athmgb15* mutants, as the role of jasmonate in stamen and pollen development has been shown previously (Huang et al., 2017b).

### AtHMGB15 regulates jasmonic acid biosynthesis and signalling during pollen development

Previous studies have established the phytohormone jasmonic acid as one of the major plant hormones required for different stages of flower development, including regulation of anther development, stamen elongation, dehiscence, flower opening and pollen development (Huang et al., 2017a, Huang et al., 2020, Huang et al., 2017b, Ishiguro et al., 2001, Mandaokar and Browse, 2009, Mandaokar et al., 2006, McConn and Browse, 1996, Qi et al., 2015). Mutants deficient in jasmonic acid biosynthesis and signalling were found to have reduced fertility or are male sterile (Cheng et al., 2009, Feys et al., 1994, Ishiguro et al., 2001, Park et al., 2002, Xie et al., 1998). JA signal should be attenuated at an appropriate period with appropriate amplitude during the development process for proper growth and fitness of the plant. This is achieved by a remarkable regulation between JA biosynthesis and JA signalling through positive and negative feedback loops (Wasternack, 2019, Wasternack and Hause, 2013, Zander et al., 2020). While positive feedback increases the jasmonate biosynthesis to activate jasmonate signalling; the negative feedback regulates the activity of TF like MYC2, by activating the expression of negative repressors like JAZ or JAZ splice variants to attenuate the JA signalling(Chini et al., 2007, Chini et al., 2016, Pauwels and Goossens, 2011, Song et al., 2011, Song et al., 2014).

### Regulation of MYC2 mediated JA signalling

The basic-helix-loop-helix transcription factor MYC2 is the master regulator of JA-signalling (Kazan and Manners, 2013). The MYC2-dependent transcription of JA responsive genes is tightly regulated by the activity of SCF^COI1^-JAZ complex. JAZ1 physically interacts with MYC2 and inhibits its transcriptional activity (Acosta and Przybyl, 2019, Chini et al., 2007, Xu et al., 2002). Jasmonate induces SCF^COI1^-dependent proteasomal degradation of JAZ and releases MYC2 for transcriptional activation (Devoto et al., 2002, Kazan and Manners, 2008, Thines et al., 2007). Interestingly, one of the primary targets of MYC2 is the promoter of *MYC2* itself along with that of *JAZ* genes during jasmonate response, indicating that MYC2 activates its transcription as well as its negative regulator, *JAZ* (Dombrecht et al., 2007, Kazan and Manners, 2013, Zander et al., 2020). Thus, JA dependent destruction of MYC2 repressor for activating JA-responsive, followed by MYC2 dependent activation of *JAZ* repressor, indicates the involvement of a negative feedback loop in JA signalling (Chini et al., 2007). Our results suggest that AtHMGB15 directly binds to the promoter region of *MYC2* gene and positively activates the transcription of *MYC2*. Since the expression of *MYC2* is compromised in AtHMGB15 deletion lines, expression of most of the *JAZ* genes (*JAZ1,5,6,7,8* and 10) were found to be down-regulated in *athmgb15* mutant (Fig. S6). Down-regulation of MYC2 and JAZ genes suggest that fine tuning of JA signalling is severely affected in *athmgb15* mutants.

### Regulation of JA biosynthesis

The expression of JA biosynthesis genes such as *DAD1, AOS, AOC1, OPR1, LOX4* and *JAR1* were significantly down-regulated in *athmgb15* mutants. Studies have shown that JA biosynthesis is regulated by a positive feedback loop through SCFCOI1-JAZ regulatory module in presence of jasmonate derivative (Devoto et al., 2002). The proteasomal degradation of JAZ repressor in presence of JA-Ile releases MYC2 to bind to JA-responsive elements (G-box) present in the promoters of JA-biosynthesis genes such as *AOS*, *AOC3, OPR3, OPRL1, LOX3* and *LOX4* to promote the transcription (Dombrecht et al., 2007, Figueroa and Browse, 2012, Kazan and Manners, 2013, Pozo et al., 2008). The MYC2 dependent transcription of JA biosynthesis genes was further supported by a previous comparative RNA-seq study showing down-regulation of JA biosynthesis genes in *MYC2* mutant *jin1-8* plants (Lorenzo et al., 2004). Additionally, our results indicate significant down-regulation of JA biosynthesis and signalling genes along with significant decrease in jasmonate content in *myc2-2*, and *jin1-2* mutants. Since the expression of *MYC2* is significantly down-regulated in *athmgb15* mutant, we have also observed low expression of JA biosynthesis genes and significant low level of jasmonate in the deletion line. Taken together, our results propose that AtHMGB15 positively regulates *MYC2* transcription for the expression of JA biosynthesis genes during pollen development.

DAD1 is a chloroplastic phospholipase A1 lipase that is involved in the initial step of JA biosynthesis for the formation of α-linonenic acid. *dad1* mutants were found to be defective in anther dehiscence, pollen maturation, and flower bud development (Ishiguro et al., 2001, Peng et al., 2013). The expression of *DAD1* is regulated by homeotic protein AGAMOUS and Auxin responsive factors ARF6 and ARF8 (Nagpal et al., 2005, Tabata et al., 2010). Our study shows no change in the expression of *AGAMOUS* or *ARFs*, however, the expression of *COI1* was found to be repressed in *athmgb15* mutants. Study has shown that wound induced expression of *DAD1* is lower in JA biosynthesis mutant *aos* and *opr3* and completely abolished in *coi1* mutant suggesting that *DAD1* expression is regulated by both COI-dependent and independent mechanisms (Ruduś et al., 2014, Hyun et al., 2008). Considering these findings, we suggest that transcription of *DAD1* is COI1-dependent during the pollen development process.

JAR1, CYP94B3 and ST2A are jasmonic acid catabolic enzymes required for the formation of jasmonic acid derivatives JA-Ile, 12-hydroxy-JA-Ile and 12-HSO4-JA respectively (Ruan et al., 2019, Wasternack, 2019). For JAR1, jasmonic acid is the substrate for JA-Ile formation; CYP94B3 use JA-Ile for hydroxylation and STA2 uses 12-OH-JA for sulphated derivate (Wasternack and Hause, 2013). Thus, it appears that the biosynthesis of JA catabolites depends upon the availability of its substrate. Since the jasmonic acid content of *athmgb15* mutant plants was found to be lower compared to wild-type, we hypothesized that the synthesis of jasmonate derivatives will be lower in mutant plants. Therefore, the expression of genes responsible for the formation of JA-derivatives will be repressed due to positive feedback. Further, Koo *et al* have demonstrated that the expression of *CYP94B3* is dependent on COI1, as the expression is completely diminished in *coi1* mutant. Therefore, reduced expression of *CYP94B3* in *athmgb15* mutant may be due to both, down-regulation of *COI1* gene expression and substrate availability (Koo et al., 2011).

### Repression of Jasmonic acid biosynthesis and signalling causes down-regulation of JA-responsive transcription factor *MYB21* and *MYB24* in *athmgb15* mutant

The R2R3-MYB transcription factors, MYB21 and MYB24, are considered as the master regulator of JA signalling during stamen development (Huang et al., 2017a, Huang et al., 2020, Yang et al., 2020). *MYB21* and *MYB24* expression were found to be down-regulated in *athmgb15* mutant. Furthermore, we have shown that AtHMGB15 binds to the promoter of *MYB24* and MYB21 and activates their transcription. There may be two possible reasons for the repression of MYBs in *athmgb15* mutant. Firstly, repression of JA biosynthesis in *athmgb15* causes down-regulation of *MYB21* and *MYB24* expression. This can be supported by a previous studies showing down-regulation of *MYB21* and *MYB24* expression (5-fold) in *opr3* mutants (Huang et al., 2020, Song et al., 2011). Also, the expression of *MYB21* and *MYB24* in *opr3* mutant can be restored by exogenous application of JA, suggesting that JA deficiency blocks the expression of these transcription factors. Additionally, Cheng et al have shown that GA-dependent expression of *DAD1*is a prerequisite for the expression of *MYB21* and *MYB24*, suggesting that GA-induced JA biosynthesis regulates the expression of *MYB21* and *MYB24*. The second possibility for the repression of MYBs in AtHMGB15 deletion lines may be due to its role as a transcription activator for the expression of *MYB21* and *MYB24*. Genetic analysis has indicated that MYB21 and MYB24 are indispensable for stamen growth and development and, *myb21myb24* double mutant is completely male sterile with short filaments, delayed anther dehiscence and non-viable pollens (Huang et al., 2020). Interestingly, overexpression of *MYB21* partially restores male sterility in *coi1* and completely restores stamen elongation and fertility in *opr3* mutant (Qi et al., 2015). As mentioned earlier, deletion of MYBs or in JA-biosynthesis mutants, stamen growth was found to be arrested so that anther fails to reach stigma for pollination. In *athmgb15* flowers, we have found that 30% of flowers showed short filaments. This may be another possible reason for having less seed yields in *athmgb15* mutant.

### AtHMGB15 interacts with MYC2 to form the activator complex for regulating JA-responsive transcription

One of the interesting findings from this study is the physical interaction of AtHMGB15 protein with MYC2 transcription factor. Our finding indicates that AtHMGB15 together with MYC2 activates the transcription of MYC2. A previous study by Zander *et al* have identified that MYC2 binds many targets that do not have canonical G-box DNA sequence motifs (Dombrecht et al., 2007, Figueroa and Browse, 2012, Kazan and Manners, 2013, Pozo et al., 2008, Zander et al., 2020). These targets have AtHMGB15 binding site as one of the enriched motifs present suggesting that MYC2 may bind indirectly to many such targets through its partner protein AtHMGB15. This study gave a clue that MYC2 needs partner protein such as AtHMGB15 for its activity. Since we have observed direct interaction of MYC2 and AtHMGB15, we believe that the interaction between these two proteins acts as a transcription activator complex in many MYC2 dependent gene expressions during JA signalling. We have observed that other than MYC2, AtHMGB15 also activate the promoters of MYB transcription factor.

In this study, we have for the first time, identified the mechanistic role of ARID/HMG group of nuclear protein in the pollen development process. We have identified the role of AtHMGB15 in the formation of pollen cell wall by positively regulating the expression of a couple of cell wall genes. We have also demonstrated how the role of AtHMGB15 in JA signalling by forming an activator complex with MYC2 transcription factor to activate JA-dependent gene expression during pollen development (Fig. 7D). To date, very less information is available regarding the physiological roles of plant ARID/HMGs, especially in gene regulation and chromatin remodelling. The present study shall be a step forward in this direction and has established a new role of AtHMGB15 in transcription activation other than being an HMG-box group of nuclear architectural protein.

## MATERIALS AND METHODS

### Plant materials and growth conditions

Arabidopsis thaliana ecotype Columbia-O (Col) was used in this study. All the mutants and over-expression lines used in this study were in the Col background. The T-DNA insertion line of AtHMGB15 (GABI_351D08) was obtained from Eurasian Arabidopsis Stock Centre (NASC). Seeds of *MYC2* mutants (*myc2-2* and *jin1-2*). The seeds were grown on Murashige and Skoog Agar plates at 22 °C under 16 h:8h light (~150 ± 10 μmol m-2 s-1) and dark cycle in the growth chamber. 20days old seedlings were transferred to soil pots in greenhouse with 60% relative humidity. Freshly opened flowers were collected every day between 9:00 am-11:00 am IST during the flowering stage (flowering stage 13) for downstream experiments. Pictures of Wildtype, *athmgb15-4* mutant, *athmgb15-4-OEA4* (RE) plants at various growth stages (rosette, inflorescence bolting, fully mature plant with flower and silique stage) were taken using a digital camera. Individual organs such as the flowers and siliques were isolated and investigated for Leica stereo-zoom microscope S9i.

#### Generation of Transgenic plants

The coding sequence of *AtHMGB15* was cloned under 35S in pMDC84 using gateway cloning system (Invitrogen). This construct was used to generate complementation lines constructed in the Col-0 and in the *athmgb15-4* mutant background. Plant transformation was performed by Agrobacterium tumefaciens-mediated floral dip method and transgenic plants were selected by hygromycin selection. The complementation lines were confirmed by PCR for insertion of the DNA fragment and qRT-PCR for expression. The list of primers used for this study is presented in supplementary table S1.

### RNA extraction, Ilumina Sequencing and q-RTPCR

Total RNA was isolated from 200mg of young flowers (flowering stage 12-13) of wildtype and *athmgb15-4* mutant using RNASure® Mini Kit (Nucleopore-Genetix). RNA isolated from three such replicates were pooled and used for illumine sequencing using 2 x 75 bp chemistry generating 30 million paired-end reads per sample. Processing of raw read, adaptor removing using Trimmomatic v0.35 and mapping of read to Arabidopsis genome (TAIR10) using TopHat v2.1.1 were performed as mentioned earlier. The differential gene expression analysis was carried out using Cufflink v1.3.0 where threshold fold Change was set (FC) values greater than zero along with P value threshold of 0.05 where threshold fold Change was set (FC) values greater than ±1 with P value cut-off filter of 0.05 were considered as differentially expressed genes. For qRTPCR was performed as described earlier. The relative fold change for the gene of interest was calculated with respect to housekeeping gene *AtEF1α* transcript (At1g07920) level using 2^-ΔΔCT^ method. The significance of the results was analysed by paired two-tailed Student’s t-test (P ≤ 0.05) using at least three independent biological replicates. The primers used in the analysis are enlisted in Supplementary Table S1.

### Bioinformatic analysis

The functional annotation clustering and KEGG pathway were generated for the significant DEGs and were analysed using DAVID v6.8. The heatmaps were generated using MeV (v4.9.0). The promoter sequence of *MYC2, MYB24, MYB21* was analysed using PlantPAN 3.0.

### Chromatin immunoprecipitation and ChIP-qPCR

Nuclei from the 700mg of wildtype flower tissue were isolated using the Plant Nuclei isolation kit (Sigma, # CELLYTPN 1) conferring to the manufacturer’s protocol. The chromatin immunoprecipitation assay was performed as described previously by (Mallik et al., 2020). and immunoprecipitated DNA was analysed by ChIP-qPCR. The data were normalized with respect to input and fold change was calculated against previously characterised two loci At1g01840 and At1g01310 using 2^-ΔΔCT^ method. Three independent biological replicate samples were used for qPCR experiments, where each sample was collected from ≥80 wildtype plants in the flowering stage. The significance of the results was analysed by paired two-tailed Student’s t-test P ≤0.05. The primer list for the ChIP study is attached in the supplementary table S1.

### DNA Binding Assay

Electrophoretic mobility shift assay EMSA was performed using the protocol described previously (Roy et al., 2016). DNA fragments (200bp) w from the promoter/upstream region of *MYC2, MYB21* and *MYB24* containing previously identified AtHMGB15 binding site ATA—(A/T)(A/T)” was PCR amplified and end-labelled with □DP^32^ATP. 5×104cpm γP32 labelled DNA (~7 fmol) was mixed with increasing concentrations of AtHMGB15 from 0.5μM to 3μM and the DNA-protein mixture was analysed by 5% native PAGE in 0.5X TBE at 4°C.

### Scanning Electron Microscopy

Pollen grains were isolated from anthers of dried flowers of wildtype, *athmgb15-4* mutant and *athmgb15-4-OEA4* (*RE*) and refined by passing them through a series of fine mesh with decreasing porosity. The pollen grains were brushed onto the brass stub with a carbon tape and subjected to gold coating in Edward gold sputter coater. The coated samples were visualised in SEM (FEI 200) under an accelerating voltage of 5, 10 and 20 kV.

### Pollen germination and viability assay

Pollen germination assay was done as described previously (Li, 2011). Pollen was isolated from mature wildtype and *athmgb15-4* flowers by drying them and then suspending them in a pollen germination medium containing 20% (w/v) Sucrose, 100mM boric acid, 1M CaCl^2^, 200mM Tris MES, 1M MgSO_4_, 30% PEG 4000 and 500mM KCl of pH 5.6-6 (Fan et al., 2001). Pollen germination was observed after 2hr, 4hr, 6hr and 24hrs and visualized by microscope (Nikon ECLIPSE Ni). Double staining with fluorescein diacetate and propidium iodide was performed using the method of Chang et al. (2014). A drop containing the stained pollens were viewed under a fluorescence microscope (Nikon ECLIPSE Ni) at 537nm and 480nm wavelengths for PI and FDA respectively.

### Hormone Estimation

Jasmonic acid content was estimated using Electron Spray Ionisation coupled with Mass Spectroscopy (ESI-MS) as described previously (Liu et al., 2010). 500mg of fresh flower tissue from wildtype, *athmgb15-4* and *athmgb15-4-OEA4 (RE*) plants were homogenised in liquid N_**2**_ and extracted overnight with Methanol (HPLC grade) at 4°C. The homogenates were centrifuged and diluted with water (HPLC grade) and subjected to the Sep-pak C18 cartridge (SPE). SPE cartridge was washed with 20% and 30% methanol and finally eluted with 100% methanol. The eluant was 10 times diluted with methanol and analysed by ESI-MS. Analytical standards of Methyl Jasmonate (Sigma® #392707), Jasmonic Acid (Sigma® #J2500) and Jasmonic Acid-Isoleucine (Cayman Chemical® #10740) were used. The relative abundance of all three derivatives in the wildtype, *athmgb15-4* and *athmgb15-4-OEA4 (RE*) samples was obtained and expressed as fold change with respect to wildtype.

### Plant treatment

For Methyl Jasmonate (MeJA) treatment, the wildtype and *athmgb15-4* plants were grown directly in soil. At the onset of flowering, 0.5 mM and 2 mM MeJA (Sigma® #392707) was sprayed directly onto the flower buds twice a day for two consecutive days. The treated flowers were harvested and used for pollen germination assay (Park et al., 2002).

### GUS Assay

GUS assay was performed as described previously (Bedi and Nag Chaudhuri, 2018). 2KB promoter regions of *MYC2, MYB21* and *MYB24* were cloned into pKGWFS7 vector, containing GUS as the reporter gene, by the Gateway cloning (Invitrogen®). Similarly, the full length coding sequence of *AtHMGB15* and *MYC2* was cloned in pMDC84 and pCambia1304 respectively. Overnight culture of *Agrobacterium tumefaciens* strain EHA105 containing *pMYC2, pMYB21, pMYB24* was mixed individually with different combinations of Agrobacterium strain containing 35S::AtHMGB15 and 35S::MYC2 at OD_600_0.8 and infiltered into the leaves of 6weeks old *Nicotiana tabacum* plants. The leaf samples after 48hrs of incubation were homogenized and GUS activity was measured using 1 mM MUG at fluorescence at 455 nm (excitation at 365 nm) in a fluorimeter (Thermo Scientific Varioskan Flash). The total protein concentration of extracted leaf samples was measured by Bradford method at 595 nm. GUS activity was represented as nanomoles of 4-MU produced per mg of protein and the total data was obtained from 15 sets of biological repeats.

### BiFC

For BiFC assay, full-length coding sequence of *AtHMGB15* and *MYC2* were cloned through Gateway cloning system (Invitrogen) into the binary vector pSITE-cEYFP-N1 (CD3-1651) and pSITE-nEYFP-C1 (CD3-1648) respectively. Agrobacterium strain (EHA105) transformed with the cloned vectors along with the empty vectors as control were infiltered into onion epidermis as done previously (Roy et al., 2019). The inner epidermal peels were isolated and subjected to wash with 1% PBS for 16 h, after which they were mounted on slide and observed for interaction under the confocal microscope.

## Data Availability

The datasets generated during this current study are available in the NCBI Sequence Read Archive repository (https://www.ncbi.nlm.nih.gov/sra/PRJNA874885) under the Accession ID: PRJNA874885.

## ACKNOWLEDGEMENTS AND FUNDING

This work was supported by SERB, Department of Science and Technology, Government of India, (SERB/2017/000768). Sonal Sachdev sincerely acknowledges UGC; Government of India, for her fellowship [UGC-Ref. No.: 749/ (CSIR-UGC NET DEC. 2016)].

The authors sincerely acknowledge Bose Institute for institutional support. The authors sincerely thanks Dr. Sreeramaiah Gangappa (IISER, Kolkata) for the seeds of SALK_083483 (*atmyc2-2*) and SALK_061267 (*atjin1-2*) and Dr. Anindita Seal (Department of Biotechnology, University of Calcutta) for BiFC vectors and Ms Ayantika Nandi for her technically support in this project. Authors would like to acknowledge Bose Institute for providing infrastructure for experiments and data analysis. Authors declared no conflict of interest.

## AUTHORS CONTRIBUTION

SC conceptualized the idea and supervised the project. SS performed experiments related to raising transgenic and morphological studies, q-RTPCR, pollen germination, EMSA, BiFC, SEM, hormone estimation and analyzed RNA seq data. RB performed experiments pollen viability, pollen germination, promoter assay, flower morphology, analyzed RNA seq data and assisted SS in EMSA, SEM and AR screened *athmgb15-4* mutant line, standardization of pollen SEM and prepared samples for RNA seq. SC wrote the original draft and all the authors read, edited and reviewed, Funding Acquisition, S.C

## REFERENCE

Acosta, I. F. & Przybyl, M. 2019. Jasmonate signaling during Arabidopsis stamen maturation. Plant and Cell Physiology, 60, 2648–2659.

An, C., Deng, L., Zhai, H., You, Y., Wu, F., Zhai, Q., Goossens, A. & Li, C. 2022. Regulation of jasmonate signaling by reversible acetylation of TOPLESS in Arabidopsis. Molecular Plant, 15, 1329–1346.

Bedi, S. & Nag Chaudhuri, R. 2018. Transcription factor ABI 3 auto-activates its own expression during dehydration stress response. FEBS letters, 592, 2594–2611.

Boter, M., Ruíz-Rivero, O., Abdeen, A. & Prat, S. 2004. Conserved MYC transcription factors play a key role in jasmonate signaling both in tomato and Arabidopsis. Genes & development, 18, 1577–1591.

Caldelari, D., Wang, G., Farmer, E. E. & Dong, X. 2011. Arabidopsis lox3 lox4 double mutants are male sterile and defective in global proliferative arrest. Plant molecular biology, 75, 25–33.

Cheng, H., Song, S., Xiao, L., Soo, H. M., Cheng, Z., Xie, D. & Peng, J. 2009. Gibberellin acts through jasmonate to control the expression of MYB21, MYB24, and MYB57 to promote stamen filament growth in Arabidopsis. PLoS genetics, 5, e1000440.

Chini, A., Boter, M. & Solano, R. 2009. Plant oxylipins: COI1/JAZs/MYC2 as the core jasmonic acid-signalling module. The FEBS journal, 276, 4682–4692.

Chini, A., Fonseca, S., Fernandez, G., Adie, B., Chico, J., Lorenzo, O., García-Casado, G., LóPez-Vidriero, I., Lozano, F. & Ponce, M. 2007. The JAZ family of repressors is the missing link in jasmonate signalling. Nature, 448, 666–671.

Chini, A., Gimenez-Ibanez, S., Goossens, A. & Solano, R. 2016. Redundancy and specificity in jasmonate signalling. Current opinion in plant biology, 33, 147–156.

Devoto, A., Nieto-Rostro, M., Xie, D., Ellis, C., Harmston, R., Patrick, E., Davis, J., Sherratt, L., Coleman, M. & Turner, J. G. 2002. COI1 links jasmonate signalling and fertility to the SCF ubiquitin–ligase complex in Arabidopsis. The Plant Journal, 32, 457–466.

Dombrecht, B., Xue, G. P., Sprague, S. J., Kirkegaard, J. A., Ross, J. J., Reid, J. B., Fitt, G. P., Sewelam, N., Schenk, P. M., Manners, J. M. & Kazan, K. 2007. MYC2 Differentially Modulates Diverse Jasmonate-Dependent Functions in Arabidopsis. The Plant Cell, 19, 2225–2245.

Feys, B. J., Benedetti, C. E., Penfold, C. N. & Turner, J. G. 1994. Arabidopsis mutants selected for resistance to the phytotoxin coronatine are male sterile, insensitive to methyl jasmonate, and resistant to a bacterial pathogen. The Plant Cell, 6, 751–759.

Figueroa, P. & Browse, J. 2012. The Arabidopsis JAZ2 promoter contains a G-Box and thymidine-rich module that are necessary and sufficient for jasmonate-dependent activation by MYC transcription factors and repression by JAZ proteins. Plant and Cell Physiology, 53, 330–343.

Gao, C., Qi, S., Liu, K., Li, D., Jin, C., Li, Z., Huang, G., Hai, J., Zhang, M. & Chen, M. 2016. MYC2, MYC3, and MYC4 function redundantly in seed storage protein accumulation in Arabidopsis. Plant Physiology and Biochemistry, 108, 63–70.

Goldberg, R. B., Beals, T. P. & Sanders, P. M. 1993. Anther development: basic principles and practical applications. The Plant Cell, 5, 1217.

Goossens, J., Mertens, J. & Goossens, A. 2017. Role and functioning of bHLH transcription factors in jasmonate signalling. Journal of Experimental Botany, 68, 1333–1347.

Hansen, F. T., Madsen, C. K., Nordland, A. M., Grasser, M., Merkle, T. & Grasser, K. D. 2008. A novel family of plant DNA-binding proteins containing both HMG-box and AT-rich interaction domains. Biochemistry, 47, 13207–13214.

Huang, H., Gao, H., Liu, B., Qi, T., Tong, J., Xiao, L., Xie, D. & Song, S. 2017a. Arabidopsis MYB24 regulates jasmonate-mediated stamen development. Frontiers in Plant Science, 8, 1525.

Huang, H., Gong, Y., Liu, B., Wu, D., Zhang, M., Xie, D. & Song, S. 2020. The DELLA proteins interact with MYB21 and MYB24 to regulate filament elongation in Arabidopsis. BMC plant biology, 20, 1–9.

Huang, H., Liu, B., Liu, L. & Song, S. 2017b. Jasmonate action in plant growth and development. Journal of experimental botany, 68, 1349–1359.

Hyun, Y., Choi, S., Hwang, H.-J., Yu, J., Nam, S.-J., Ko, J., Park, J.-Y., Seo, Y. S., Kim, E. Y. & Ryu, S. B. 2008. Cooperation and functional diversification of two closely related galactolipase genes for jasmonate biosynthesis. Developmental cell, 14, 183–192.

Ishiguro, S., Kawai-Oda, A., Ueda, J., Nishida, I. & Okada, K. 2001. The DEFECTIVE IN ANTHER DEHISCENCE1 Gene Encodes a Novel Phospholipase A1 Catalyzing the Initial Step of Jasmonic Acid Biosynthesis, Which Synchronizes Pollen Maturation, Anther Dehiscence, and Flower Opening in Arabidopsis. The Plant Cell, 13, 2191–2209.

Jang, G., Yoon, Y. & Choi, Y. D. 2020. Crosstalk with jasmonic acid integrates multiple responses in plant development. International journal of molecular sciences, 21, 305.

Kazan, K. & Manners, J. M. 2008. Jasmonate signaling: toward an integrated view. Plant physiology, 146, 1459–1468.

Kazan, K. & Manners, J. M. 2013. MYC2: the master in action. Molecular plant, 6, 686–703.

Koo, A. J., Cooke, T. F. & Howe, G. A. 2011. Cytochrome P450 CYP94B3 mediates catabolism and inactivation of the plant hormone jasmonoyl-L-isoleucine. Proceedings of the National Academy of Sciences, 108, 9298–9303.

Li, X. 2011. Arabidopsis pollen tube germination. Bio-protocol, e73–e73.

Liu, X., Yang, Y., Lin, W., Tong, J., Huang, Z. & Xiao, L. 2010. Determination of both jasmonic acid and methyl jasmonate in plant samples by liquid chromatography tandem mass spectrometry. Chinese Science Bulletin, 55, 2231–2235.

Lorenzo, O., Chico, J. M., Saénchez-Serrano, J. J. & Solano, R. 2004. JASMONATE-INSENSITIVE1 Encodes a MYC Transcription Factor Essential to Discriminate between Different Jasmonate-Regulated Defense Responses in Arabidopsis[W]. The Plant Cell, 16, 1938–1950.

Mallik, R., Prasad, P., Kundu, A., Sachdev, S., Biswas, R., Dutta, A., Roy, A., Mukhopadhyay, J., Bag, S. K. & Chaudhuri, S. 2020. Identification of genome-wide targets and DNA recognition sequence of the Arabidopsis HMG-box protein AtHMGB15 during cold stress response. Biochimica et Biophysica Acta (BBA)-Gene Regulatory Mechanisms, 1863, 194644.

Mandaokar, A. & Browse, J. 2009. MYB108 acts together with MYB24 to regulate jasmonate-mediated stamen maturation in Arabidopsis. Plant Physiology, 149, 851–862.

Mandaokar, A., Thines, B., Shin, B., Markus Lange, B., Choi, G., Koo, Y. J., Yoo, Y. J., Choi, Y. D., Choi, G. & Browse, J. 2006. Transcriptional regulators of stamen development in Arabidopsis identified by transcriptional profiling. The Plant Journal, 46, 984–1008.

Marciniak, K. & Przedniczek, K. 2019. Comprehensive Insight into Gibberellin- and Jasmonate-Mediated Stamen Development. Genes, 10, 811.

Mascarenhas, J. P. 1990. Gene activity during pollen development. Annual review of plant biology, 41, 317–338.

Mcconn, M. & Browse, J. 1996. The Critical Requirement for Linolenic Acid Is Pollen Development, Not Photosynthesis, in an Arabidopsis Mutant. The Plant Cell, 8, 403–416.

Mccormick, S. 2004. Control of Male Gametophyte Development. The Plant Cell, 16, S142–S153.

Nagpal, P., Ellis, C. M., Weber, H., Ploense, S. E., Barkawi, L. S., Guilfoyle, T. J., Hagen, G., Alonso, J. M., Cohen, J. D. & Farmer, E. E. 2005. Auxin response factors ARF6 and ARF8 promote jasmonic acid production and flower maturation.

Park, J. H., Halitschke, R., Kim, H. B., Baldwin, I. T., Feldmann, K. A. & Feyereisen, R. 2002. A knock-out mutation in allene oxide synthase results in male sterility and defective wound signal transduction in Arabidopsis due to a block in jasmonic acid biosynthesis. The Plant Journal, 31, 1–12.

Pauwels, L. & Goossens, A. 2011. The JAZ proteins: a crucial interface in the jasmonate signaling cascade. The Plant Cell, 23, 3089–3100.

Peng, Y. J., Shih, C. F., Yang, J. Y., Tan, C. M., Hsu, W. H., Huang, Y. P., Liao, P. C. & Yang, C. H. 2013. A RING-type E 3 ligase controls anther dehiscence by activating the jasmonate biosynthetic pathway gene DEFECTIVE IN ANTHER DEHISCENCE 1 in A rabidopsis. The Plant Journal, 74, 310–327.

Pozo, M. J., Van Der Ent, S., Van Loon, L. & Pieterse, C. M. 2008. Transcription factor MYC2 is involved in priming for enhanced defense during rhizobacteria-induced systemic resistance in Arabidopsis thaliana. New phytologist, 180, 511–523.

Qi, T., Huang, H., Song, S. & Xie, D. 2015. Regulation of Jasmonate-Mediated Stamen Development and Seed Production by a bHLH-MYB Complex in Arabidopsis. The Plant Cell, 27, 1620–1633.

Roy, A., Dutta, A., Roy, D., Ganguly, P., Ghosh, R., Kar, R. K., Bhunia, A., Mukhopadhyay, J. & Chaudhuri, S. 2016. Deciphering the role of the AT-rich interaction domain and the HMG-box domain of ARID-HMG proteins of Arabidopsis thaliana. Plant Mol Biol, 92, 371–88.

Ruan, J., Zhou, Y., Zhou, M., Yan, J., Khurshid, M., Weng, W., Cheng, J. & Zhang, K. 2019. Jasmonic Acid Signaling Pathway in Plants. International Journal of Molecular Sciences, 20, 2479.

Ruduś, I., Terai, H., Shimizu, T., Kojima, H., Hattori, K., Nishimori, Y., Tsukagoshi, H., Kamiya, Y., Seo, M. & Nakamura, K. 2014. Wound-induced expression of DEFECTIVE IN ANTHER DEHISCENCE1 and DAD1-like lipase genes is mediated by both CORONATINE INSENSITIVE1-dependent and independent pathways in Arabidopsis thaliana. Plant cell reports, 33, 849–860.

Sanders, P. M., Bui, A. Q., Weterings, K., Mcintire, K., Hsu, Y.-C., Lee, P. Y., Truong, M. T., Beals, T. & Goldberg, R. 1999. Anther developmental defects in Arabidopsis thaliana male-sterile mutants. Sexual plant reproduction, 11, 297–322.

Schweizer, F., Fernández-Calvo, P., Zander, M., Diez-Diaz, M., Fonseca, S., Glauser, G., Lewsey, M. G., Ecker, J. R., Solano, R. & Reymond, P. 2013. Arabidopsis Basic Helix-Loop-Helix Transcription Factors MYC2, MYC3, and MYC4 Regulate Glucosinolate Biosynthesis, Insect Performance, and Feeding Behavior The Plant Cell, 25, 3117–3132.

Scott, R. J., Spielman, M. & Dickinson, H. 2004. Stamen structure and function. The plant cell, 16, S46–S60.

Song, S., Qi, T., Huang, H., Ren, Q., Wu, D., Chang, C., Peng, W., Liu, Y., Peng, J. & Xie, D. 2011. The jasmonate-ZIM domain proteins interact with the R2R3-MYB transcription factors MYB21 and MYB24 to affect jasmonate-regulated stamen development in Arabidopsis. The Plant Cell, 23, 1000–1013.

Song, S., Qi, T., Wasternack, C. & Xie, D. 2014. Jasmonate signaling and crosstalk with gibberellin and ethylene. Current Opinion in Plant Biology, 21, 112–119.

Stintzi, A. & Browse, J. 2000. The Arabidopsis male-sterile mutant, opr3, lacks the 12-oxophytodienoic acid reductase required for jasmonate synthesis. Proceedings of the National Academy of Sciences, 97, 10625–10630.

ŠTros, M., Launholt, D. & Grasser, K. D. 2007. The HMG-box: a versatile protein domain occurring in a wide variety of DNA-binding proteins. Cellular and Molecular Life Sciences, 64, 2590.

Tabata, R., Ikezaki, M., Fujibe, T., Aida, M., Tian, C.-E., Ueno, Y., Yamamoto, K. T., Machida, Y., Nakamura, K. & Ishiguro, S. 2010. Arabidopsis auxin response factor6 and 8 regulate jasmonic acid biosynthesis and floral organ development via repression of class 1 KNOX genes. Plant and Cell Physiology, 51, 164–175.

Thines, B., Katsir, L., Melotto, M., Niu, Y., Mandaokar, A., Liu, G., Nomura, K., He, S. & Howe, G. 2007. Browse J. 2007. JAZ repressor proteins are targets of the SCF (COI1) complex during jasmonate signalling. Nature, 448, 661–665.

Vera-Sirera, F., Gomez, M. D. & Perez-Amador, M. A. 2016. DELLA proteins, a group of GRAS transcription regulators that mediate gibberellin signaling. Plant transcription factors. Elsevier.

Wasternack, C. 2019. Termination in jasmonate signaling by MYC2 and MTBs. Trends in plant science, 24, 667–669.

Wasternack, C. & Hause, B. 2013. Jasmonates: biosynthesis, perception, signal transduction and action in plant stress response, growth and development. An update to the 2007 review in Annals of Botany. Annals of botany, 111, 1021–1058.

Wilson, Z. A. & Zhang, D.-B. 2009. From Arabidopsis to rice: pathways in pollen development. Journal of Experimental Botany, 60, 1479–1492.

Xia, C., Wang, Y. J., Liang, Y., Niu, Q. K., Tan, X. Y., Chu, L. C., Chen, L. Q., Zhang, X. Q. & Ye, D. 2014. The ARID-HMG DNA-binding protein A t HMGB 15 is required for pollen tube growth in Arabidopsis thaliana. The Plant Journal, 79, 741–756.

Xie, D.-X., Feys, B. F., James, S., Nieto-Rostro, M. & Turner, J. G. 1998. COI1: an Arabidopsis gene required for jasmonate-regulated defense and fertility. Science, 280, 1091–1094.

Xu, L., Liu, F., Lechner, E., Genschik, P., Crosby, W. L., Ma, H., Peng, W., Huang, D. & Xie, D. 2002. The SCFCOI1 ubiquitin-ligase complexes are required for jasmonate response in Arabidopsis. The Plant Cell, 14, 1919–1935.

Yang, Z., Li, Y., Gao, F., Jin, W., Li, S., Kimani, S., Yang, S., Bao, T., Gao, X. & Wang, L. 2020. MYB21 interacts with MYC2 to control the expression of terpene synthase genes in flowers of Freesia hybrida and Arabidopsis thaliana. Journal of Experimental Botany, 71, 4140–4158.

Zander, M., Lewsey, M. G., Clark, N. M., Yin, L., Bartlett, A., Saldierna Guzmán, J. P., Hann, E., Langford, A. E., Jow, B. & Wise, A. 2020. Integrated multi-omics framework of the plant response to jasmonic acid. Nature plants, 6, 290–302.

Zhai, Q., Zhang, X., Wu, F., Feng, H., Deng, L., Xu, L., Zhang, M., Wang, Q. & Li, C. 2015. Transcriptional mechanism of jasmonate receptor COI1-mediated delay of flowering time in Arabidopsis. The Plant Cell, 27, 2814–2828.

Zhang, X., He, Y., Li, L., Liu, H. & Hong, G. 2021. Involvement of the R2R3-MYB transcription factor MYB21 and its homologs in regulating flavonol accumulation in Arabidopsis stamen. Journal of experimental botany, 72, 4319–4332.

Zhang, Z. B., Zhu, J., Gao, J. F., Wang, C., Li, H., Li, H., Zhang, H. Q., Zhang, S., Wang, D. M. & Wang, Q. X. 2007. Transcription factor AtMYB103 is required for anther development by regulating tapetum development, callose dissolution and exine formation in Arabidopsis. The Plant Journal, 52, 528–538.

